# Spatiotemporal cytokinin signaling imaging reveals IPT3 function in nodule development in *Medicago truncatula*

**DOI:** 10.1101/2021.04.23.441163

**Authors:** Paolo M. Triozzi, Thomas B. Irving, Henry W. Schmidt, Zachary P. Keyser, Sanhita Chakraborty, Kelly M. Balmant, Wendell J. Pereira, Christopher Dervinis, Kirankumar S. Mysore, Jiangqi Wen, Jean-Michel Ané, Matias Kirst, Daniel Conde

**Affiliations:** School of Forest, Fisheries and Geomatics Sciences, University of Florida, Gainesville, FL 32611, USA; Department of Bacteriology, University of Wisconsin-Madison, Madison, WI 53706, USA; Noble Research Institute, Ardmore, OK 73401, USA; Department of Agronomy, University of Wisconsin-Madison, Madison, WI 53706, USA; Genetics Institute, University of Florida, Gainesville, FL 32611, USA

**Keywords:** Cytokinin, *IPT3*, *Medicago truncatula*, *Sinorhizobium meliloti*, nodule development, nodule organogenesis, cytokinin sensor, cytokinin signaling, cytokinin biosynthesis

## Abstract

Most legumes can establish a symbiotic association with soil rhizobia that triggers the development of root nodules. These nodules host the rhizobia and allow them to fix nitrogen efficiently. The perception of bacterial lipo-chitooligosaccharide (LCO) signal in the epidermis initiates a signaling cascade that allows rhizobial intracellular infection in the root and de-differentiation and activation of cell division that gives rise to the nodule. Nodule organogenesis and rhizobial infection need to be coupled in space and time for successful nodulation. The plant hormone cytokinin (CK) acts as an essential positive regulator of nodule organogenesis, and specific CK receptors are required for nodule formation. Temporal regulation of tissue-specific CK signaling and biosynthesis in response to LCOs or *Sinorhizobium meliloti* inoculation in *Medicago truncatula* remains poorly understood. In the present study, using a fluorescence-based CK sensor (*TCSn::nls:tGFP*), we performed a high-resolution tissue-specific temporal characterization of the CK response’s sequential activation during root infection and nodule development in *M. truncatula* after inoculation with *S. meliloti*. Loss-of-function mutants of the CK-biosynthetic gene *ISOPENTENYL TRANSFERASE 3* (*IPT3*) showed impairment of nodulation, suggesting that IPT3 is required for nodule development in *M. truncatula*. Simultaneous live imaging of *pIPT3::tdTOMATO* and the CK sensor showed that *IPT3* induction in the root stele at the base of nodule primordium contributes to CK biosynthesis, which in turn promotes expression of positive regulators of nodule organogenesis in *M. truncatula*.

**One-sentence summary:** High-resolution spatiotemporal imaging of cytokinin signaling reveals IPT3 function during indeterminate nodule development in *Medicago truncatula*

## INTRODUCTION

Legume species acquired the capacity to interact symbiotically with rhizobium bacteria to fix atmospheric dinitrogen, allowing their growth without fertilizers on nitrogen-deprived soils. This interaction involves the development of specific organs, the root nodules, that host rhizobia and provide them with carbon sources and the microenvironment required for nitrogen fixation. In most rhizobia-legume associations, the perception of bacterial lipo-chitooligosaccharide (LCO) signals commonly known as Nod factors in the epidermis initiates a molecular cascade that is transmitted to the inner cell layers activating cell division, with simultaneous rhizobial infection of the host root. In *Medicago truncatula*, rhizobial infection is initiated within a curled root hair tip and the subsequent formation of a transcellular apoplastic compartment called the infection thread (IT). The IT traverses the epidermis, cortex and ramifies within the confines of the nodule primordium, developed by organized cell divisions in the root endodermis, cortex, and pericycle (Roy et al., 2020). These coordinated mechanisms of root infection and nodule organogenesis ensure that nodule maturation occurs in perfect coordination with nodule colonization by rhizobia (Xiao et al., 2014). *M. truncatula* produces indeterminate nodules that are characterized by a longitudinal gradient of differentiation with a persistent distal apical meristem and older proximal layers (Ferguson et al., 2010). Like other developmental processes, nodulation is modulated by phytohormones (Buhian and Bensmihen, 2018).

The plant hormone cytokinin (CK) is involved in various aspects of plant growth and development. CK signaling consists of a phosphorelay mediated by a two-component system, comprising a sensor and a response regulator. The site of CK binding is suggested to be the lumen of the endoplasmic reticulum. The CK-induced phosphorelay causes transcriptional changes in the nucleus mediated by type-B and type-A response regulators (RRs), which play positive, and negative roles in this regulation, respectively (Kieber and Schaller, 2018). Type-B RRs typically bind to target genes at the consensus sequence (A/G)GAT(T/C) enriched in their *cis*-regulatory regions, and synthetic CK sensors called Two-Component signaling Sensors (TCS) containing concatemeric versions of this sequence have been studied in plants (Zürcher et al., 2013). CK plays essential roles during nodule formation (Ferguson and Mathesius, 2014; Gamas et al., 2017). In *M. truncatula*, CK accumulates in the root susceptibility zone as early as 3 hours after LCO treatment (van Zeijl et al., 2015). The TCS is activated by rhizobium in the cortical cells in *M. truncatula* that forms indeterminate nodules and in *Lotus japonicus* that forms determinate ones (Held et al., 2014; Jardinaud et al., 2016). In *Glycine max*, a regulatory feedback loop involving auxin and cytokinin governs proper determinate nodule development (Turner et al., 2013). Rhizobia also induce the expression of CK biosynthetic and signaling genes in the epidermis, based on transcriptomic studies focused on the epidermal cells of *M. truncatula* (Liu et al., 2015; Damiani et al., 2016; Jardinaud et al., 2016). The *pMtRR9::GUS* transcriptional reporter, a cytokinin RR type-A (RRA), was rapidly detected in the root epidermis, in addition to other root tissues, in response to LCOs (Op den Camp et al., 2011). A more recently developed CK signaling sensor termed TCS new (TCSn) (Zürcher et al., 2013), driving GUS expression, enabled detection of the activation of a CK response in the *M. truncatula* root epidermis and the outer cortex 8 hours after the LCO treatment (Jardinaud et al., 2016). In *L. japonicus*, the *TCS* reporter was activated first in the cortex and only later in the epidermis by rhizobia (Held et al., 2014; Reid et al., 2017). The sequence of activation of CK responses during early symbiotic stages may differ between these two nodulating species, likely as a reflection of differences in the process of nodule development between them (Gamas et al., 2017).

CK plays an antagonistic role during root infection at the epidermis and nodule formation in the cortex (Gamas et al., 2017). The positive regulation of CK on nodule formation was first reported by physiological studies, which showed that exogenous CK induces the formation of nodule-like structures on the roots of several legumes (Heckmann et al., 2011). Further evidence for the positive role of CK in nodule inception has come from the analysis of nodulation-defective mutants altered in CK receptors, LOTUS HISTIDINE KINASE 1 (LHK1) in *L. japonicus*, and CYTOKININ RESPONSE 1 (CRE1) in *M. truncatula* (Gonzalez-Rizzo et al., 2006; Murray et al., 2007; Tirichine et al., 2007; Plet et al., 2011), and other CK receptors in both legumes (Held et al., 2014; Boivin et al., 2016). Moreover, gain-of-function *LHK1* and *MtCRE1* lines generate spontaneous nodules in the absence of the rhizobia (Tirichine et al., 2007; Madsen et al., 2010; Ovchinnikova et al., 2011). Transcriptomic analyses in *M. truncatula* also enabled identifying symbiotic genes that are rapidly induced by an exogenous CK on roots (Ariel et al., 2012), such as *NODULE INCEPTION* (*NIN*). NIN and the CRE1-dependent pathways are connected by a positive feedback loop, with NIN binding to the CRE1 promoter and activating its expression (Vernié et al., 2015). Similarly, *CRE1* is required for cytokinin-induced *NIN* expression (Plet et al., 2011). In *L. japonicus* roots, exogenous CK treatment also induces *NIN* specifically in root cortical cells (Heckmann et al., 2011). Moreover, *NIN* ectopic expression leads to root cortical cell divisions and nodule-like structures in both *L. japonicus* and *M. truncatula* (Soyano et al., 2013; Vernié et al., 2015), through the activation of transcription factors, such as Nuclear Factor Y subunit A1 (NFYA1) and Nuclear Factor Y subunit B1 (NFYB1) (Soyano et al., 2013; Laloum et al., 2014; Hossain et al., 2016; Shrestha et al., 2021). In contrast, in the epidermis of *M. truncatula*, CK negatively regulates root infection (Gamas et al., 2017). The epidermal CK pool was depleted by expressing a CK OXIDASE/DEHYDROGENASE (CKX) enzyme under an epidermis-specific promoter, which lead to an increased number of ITs and nodules (Jardinaud et al., 2016). Additionally, exogenous CK treatment inhibited the induction of the LCO response and pre-infection marker *MtENOD11*, in a MtCRE1 dependent fashion (Jardinaud et al., 2016). Recently, a link between the epidermis-derived CK and cortical cell divisions was established (Jarzyniak et al., 2021). *M. truncatula* ATP-binding cassette (ABC) transporter 56 (MtABCG56), transports CK from the epidermal to cortical cells, activating the CRE-dependent CK responses, including the *RRA4* (Jarzyniak et al., 2021). These downstream responses trigger further CK biosynthesis required for nodule development (Mortier et al., 2014; van; Zeijl et al., 2015; Vernié et al., 2015).

Several CK biosynthesis genes, including *ISOPENTENYL TRANSFERASE 3* (*IPT3*) and *1* (*IPT1*), *CYP735A1, LONELY GUY 1* (*LOG1*), and *2* (*LOG2*), are upregulated in response to LCOs or during nodulation in *L. japonicus* and *M. truncatula* (Chen et al., 2014; Mortier et al., 2014; Azarakhsh et al., 2015; van Zeijl et al., 2015; Azarakhsh et al., 2018; Schiessl et al., 2019). The expression of *MtIPT3, MtLOG1* and *MtLOG2* transcriptional GUS reporters were also detected in the nodule primordium (Mortier et al., 2014; Azarakhsh et al., 2020). Decreasing *LOG1* expression leads to impaired nodulation in *M. truncatula*, being involved in the nodule primordium development (Mortier et al., 2014). All these studies highlight the importance of CK biosynthesis during root infection and nodule development. Transcriptional fusions using GUS gene reporter allowed identifying CK signaling at a tissue-specific level in *M. truncatula* roots during these biological processes. However, these studies have been limited in their temporal resolution. A detailed spatial and temporal characterization of the CK response in *M. truncatula* roots should clarify the role of this hormone in nodule induction and organogenesis.

In the current study, we present the spatiotemporal regulation of CK response in rhizobia-inoculated roots using a new fluorescence-based CK signaling sensor, *pTCSn::nls:tGFP*. To further explore the potential of this sensor, we employed it along with the transcriptional fusion of the CK biosynthetic gene *IPT3* during nodule development. Simultaneous monitoring of the *pIPT3::tdTOMATO* reporter and the CK sensor activities during nodule development suggested that *IPT3* induction at the base of nodule primordium contributes to CK biosynthesis, which in turn, promotes nodule organogenesis in *M. truncatula*. Furthermore, we analyzed the loss-of-function mutant of *ipt3* and found that it is required for nodule development in *M. truncatula*.

## RESULTS

### A fluorescent protein-based cytokinin sensor is activated in root epidermal and cortical cells upon CK treatment in *Medicago truncatula*

CK responses have been studied in response to LCOs and *S. meliloti* in *M. truncatula* roots, using transcriptional reporters with *RRAs* or the synthetic *TCSn* promoters fused to the GUS gene (Op den Camp et al., 2011; Plet et al., 2011; Jardinaud et al., 2016; Fonouni-Farde et al., 2017). In soybean, fluorescent protein-based auxin and CK transcriptional reporters have been successfully used to monitor and determine their cellular level ratios in root and nodule tissues (Fisher et al., 2018). In *L. japonicus*, a tissue-specific time course experiment following the activity of *TCSn::nls:GFP* showed that CK response occurs in cortical cells before spreading to the epidermis (Reid et al., 2017).

In the present work, we designed a fluorescent protein-based CK transcriptional reporter that addresses the limitations of the GUS reporter system. This CK sensor consists of the *TCSn* promoter (Zürcher et al., 2013) driving the expression of the turbo green fluorescent protein (tGFP) fused to a nuclear localization signal peptide (*pTCSn::nls:tGFP*). This alternative approach to the GUS reporter system allows continuous, non-destructive monitoring of CK signaling throughout plant development by live imaging, and co-imaging with other fluorescence-based reporters.

Before evaluating the CK sensor activity, we characterized the timing of CK transcriptional responses in *M. truncatula* roots by analyzing the expression profiles of three *RRA* genes, *RRA3, RRA4*, and *RRA11* after 1, 8, 24, and 48 hours of 1 μM 6-benzylamino-purine (6-BAP) treatment. We found that *RRAs* reached their maximum gene expression after 24 hours of the 6-BAP treatment **(Fig. 1A)**. Based on the CK signaling activation timing, we assessed the tissue-specific CK response using the *pTCSn::nls:tGFP* CK sensor in *M. truncatula* transgenic roots after 24 hours of BAP treatment. In non-treated roots, tGFP was primarily detected in the columella, root apical meristem, and elongation zone of the root tip, as previously described for the *pTCSn::GUS* reporter (**Fig. 1B**; Jardinaud et al., 2016; Fonouni-Farde et al., 2017). Very few cells showed nuclei-localized fluorescence in the differentiation zone (DZ) of the root **(Fig. 1C)**, indicating the absence of CK response in the DZ in non-treated roots. In contrast, roots treated with 1 μM 6-BAP for 24 hours exhibited a strong induction of nuclei-localized fluorescence in the epidermis and cortex in the DZ and the differentiated zone of the root **(Fig. 1 D-E)**. DAPI counterstain confirmed that tGFP fluorescence was localized to the nuclei **(Fig. S1)**. These observations indicate that the *pTCSn::nls:tGFP* sensor constitutes a suitable molecular tool to investigate the CK response in *M. truncatula* and that CK signal transduction occurs in *M. truncatula* epidermal and cortical cells after the application of CK to the root.

**Figure 1.**
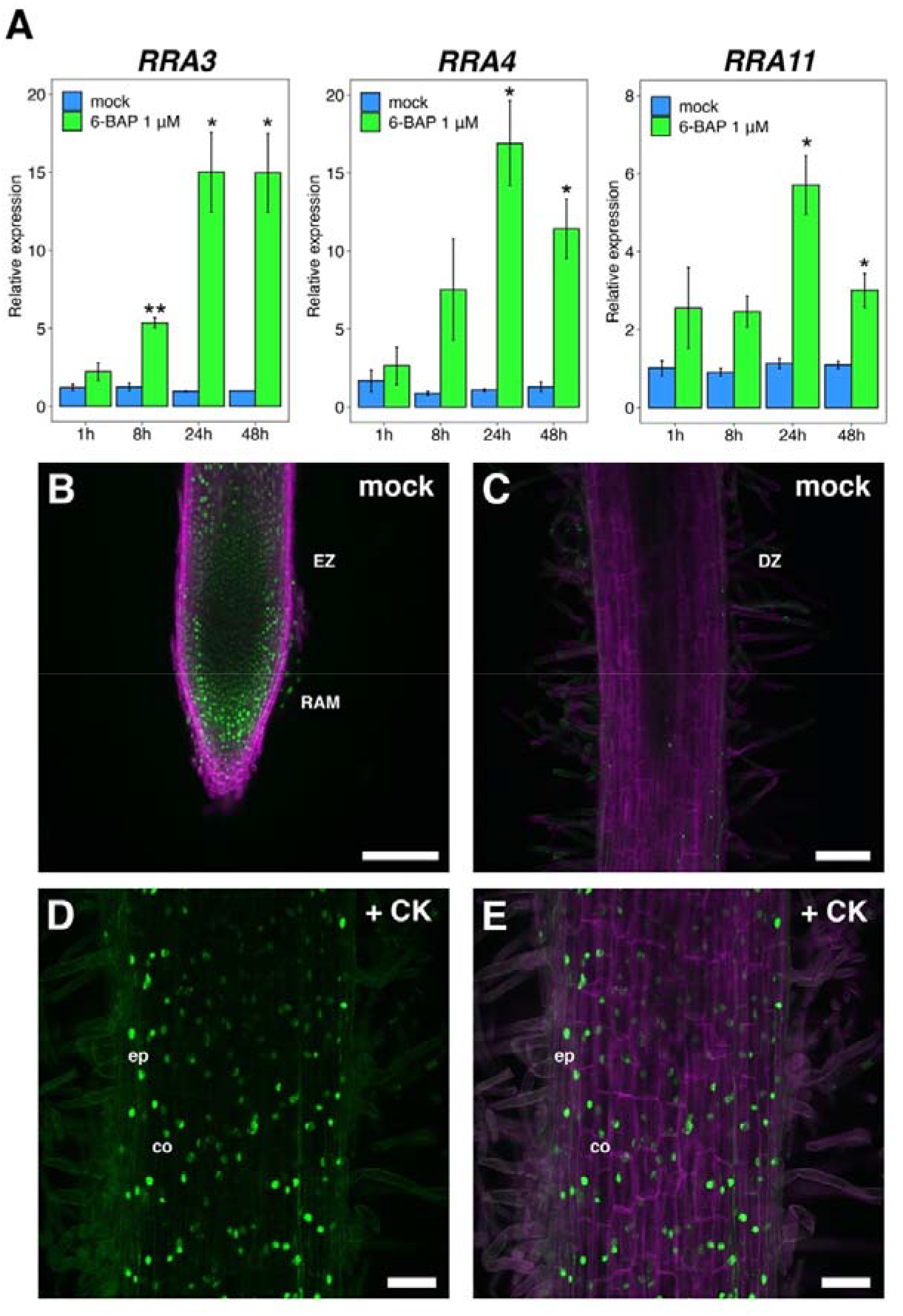
**A** reporter of CK signaling based on the *pTCSn::nls:tGFP* transcriptional fusion is activated in the epidermal and cortical cells after CK treatment. (A) qRT-PCR analyses of RRAs gene expression after 1, 8, 24, and 48 hours of 6-BAP or mock treatment. The Student’s *t*-test was performed, and asterisks represent statistically significant differences between 6-BAP and mock treatments in each time point. \**P*<0.05, \*\**P*<0.01. Values are the means ± SE of two biological replicates (n=2). (B)*pTCSn::nls:tGFP* activity in the root tip of non-treated *M, truncatula* transgenic root. The tGFP signal from the nuclei of the root apical meristem (RAM) and elongation zone (EZ) is shown in green. In magenta, the signal emitted by cell wall polysaccharides bound to Calcofluor white M2R is shown. (C) *pTCSn::nls:tGFP* activity at the differentiation zone (DZ) of non-treated transgenic root and (D, E) in DZ of 1 μM 6-BAP treated transgenic root after 24 hours in the epidermis (epi) and cortex (co). Scale bar: 100 μm in B, C and 50 μm in D, E.

### Tissue-specific time-course of CK signaling during the early symbiotic interaction with *S. meliloti* in *M. truncatula* roots

To characterize the spatial-temporal regulation of CK response during indeterminate nodule development in *M. truncatula*, we analyzed the activity of *pTCSn::nls:tGFP* in a time-course experiment using transgenic roots after *S. meliloti* inoculation. Prior to inoculation, very low *pTCSn::nls:tGFP* activity was detected in the cell layers of the susceptibility zone (SZ; **Fig. 2A**). At 4 hours after inoculation (hai), *pTCSn::nls:tGFP* activity started in the epidermal cells of the SZ, indicating that rhizobia-induced CK response in the epidermis is a very early response in *M. truncatula* **(Fig. 2B)**. At 24 hai, nuclei-localized fluorescence was still observed in the epidermal cells but was also present in outer cortical cells of the SZ **(Fig. 2C)** and by 48 hai, a strong fluorescence was widespread in the outer and inner cortical cell layers of the SZ **(Fig. 2D)**. Thus, CK signaling is activated first in the epidermis, reaches the outer cortical cells within 24 hai and extends to the majority of cortical cell layers within 48 hai.

**Figure 2.**
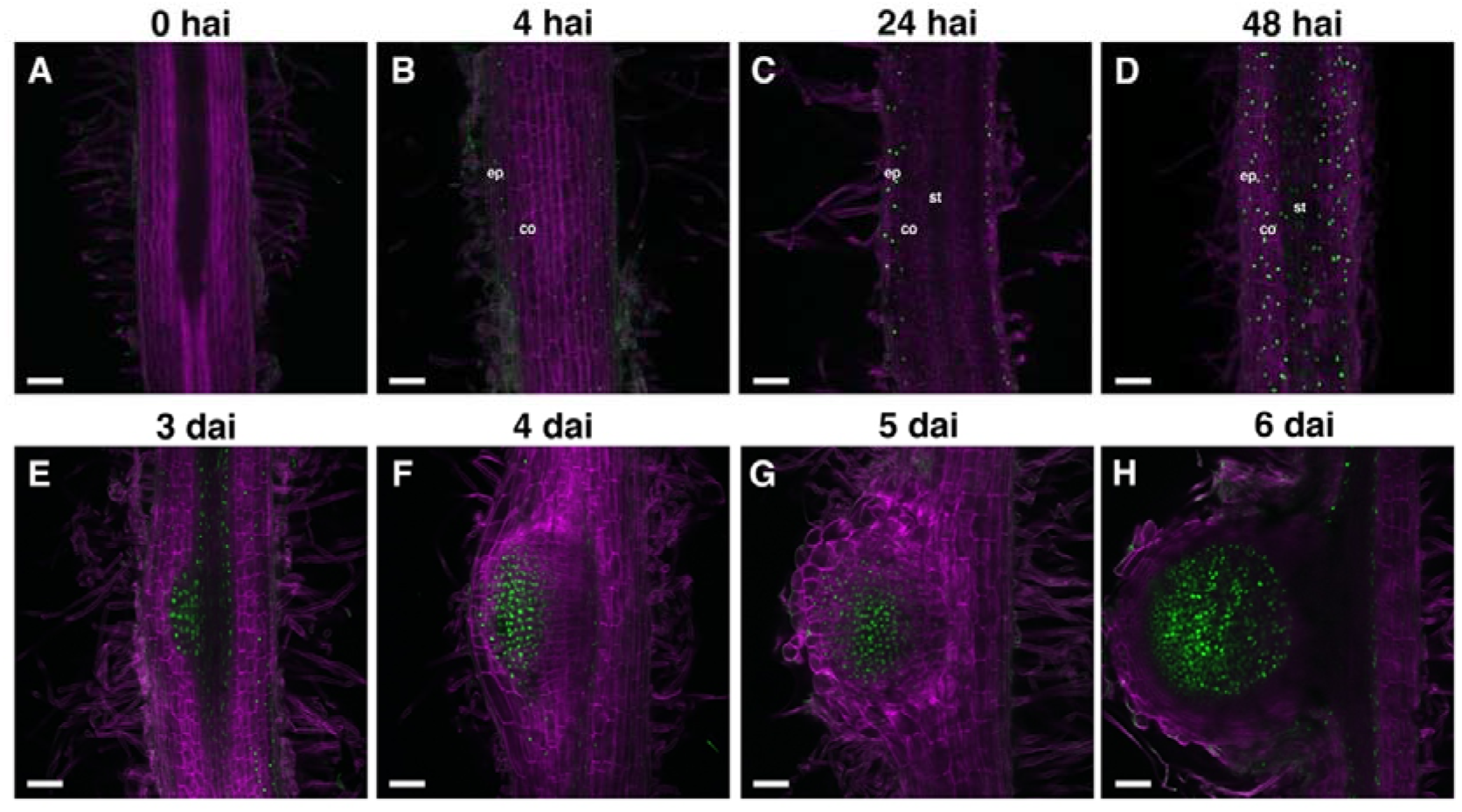
Spatio-temporal activation of cytokinin signaling during indeterminate nodule development in *M. truncatula*. (A-D) *pTCSn::nls:tGFP* activity (green) and cell walls (calcofluor white stained, magenta) in the susceptible zone of transgenic root after 0, 4, 24, and 48 hai with *S. meliloti*. Epidermis (ep), cortex (co) and stele (st). (E-H)*pTCSn::nls:tGFP* activity during nodule primordium development at (E) stages II/III, (F) stages IV/V, (G) stage VI and (H) mature nodule. Scale bar: 100 μm.

### Tissue-specific time-course of the activation of CK signaling during the indeterminate nodule development in *M. truncatula*

Characterization of CK response during indeterminate nodule development remains limited to a few time points during the process by using *RRA* promoters fused to GUS reporters in *M. truncatula* (Op den Camp et al., 2011; Plet et al., 2011). To obtain a better spatio-temporal resolution of CK response during the indeterminate nodule initiation and development, we monitored the *pTCSn::nls:tGFP* activity throughout nodule development from the first cell divisions in the pericycle, the endodermis, and the cortex, until the mature nodule formation. Moreover, we associate the tissue-specific *pTCSn::nls:tGFP* activity time course with the sequential cell division program characterized previously during nodule formation (Xiao et al., 2014).

At 3 days after inoculation (dai), the *pTCSn::nls:tGFP* signal that was widely distributed across the cortical cell layers of the SZ (see previous section) disappears, giving rise to a robust and more localized signal at the pericycle and dividing cortical cell layers C3-C5 that are related with the nodule primordium initiation, at the developmental stages II and III (**Fig. 2E**; Xiao et al., 2014). This nodule primordium-specific pattern of the CK response allowed us to clearly distinguish a nodule primordium from a lateral root primordium (LRP) during their early developmental stages, where tGFP expression was weaker and limited to the vasculature and developing meristem (**Fig. S2**). At 4 dai, we found that the stage IV and V nodule primordia showed the CK signaling activation extending to most of the dividing cortical cell layers (**Fig. 2F**). At 5 dai, the nodule primordium emerges from the main root and becomes a true nodule when the meristem starts functioning (stage VI). At this point, the CK response was localized to the C3, and the C4-C5 derived cells that form the multi-layered nodule meristem (C3) and the non-meristem zone immediately below (C4-C5), respectively **(Fig. 2G)**. At 6 dai, the nodule is in an advanced developmental stage, with the vascular bundles starting to surround the nodule meristem. At this stage, the CK signaling is strongly activated in the central zone of the nodule, including the nodule meristem and the C4/5- derived cells that will be colonized by rhizobia (**Fig. 2H**).

### Cytokinin biosynthesis by IPT3 is required for nodule development in *M. truncatula*

CK and its downstream responses are critical regulators of nodule initiation and development. However, the molecular mechanism of the local CK biosynthesis during nodule organogenesis remains poorly characterized in *M. truncatula*. It has been proposed that a KNOX3 controls the transcription of two CK biosynthesis genes, *IPT3* and *LOG2*, to promote CK biosynthesis during nodule organogenesis (Azarakhsh et al., 2015; Azarakhsh et al., 2020). However, the genetic characterization of CK biosynthesis genes in *M. truncatula* remains limited to *LOG1* (Mortier et al., 2014). Recently, it has been proposed that ABCG56 actively exports bioactive CKs from the epidermis to the cortex, driving the CRE1-dependent cortical CK response. This would lead to *de novo* CK biosynthesis in the cortex, required for nodule organogenesis (Jarzyniak et al., 2021).

It was previously shown that the *IPT3* expression is induced at 72 hai, reaching a maximum at 5 dai (Schiessl et al., 2019). Moreover, the *pIPT3::GUS* reporter showed that *IPT3* is expressed in the nodule primordium of *M. truncatula* (Azarakhsh et al., 2020). These observations indicate that IPT3 represents an excellent candidate to investigate the role of CK biosynthesis during nodule organogenesis.

We identified lines from the Noble Research Institute *Medicago truncatula Tnt1* collection showing an insertion in the single exon of *IPT3* (Tadege et al., 2008). These mutants were named *ipt3-1; ipt3-2*, and *ipt3-3* (**Fig. 3A**). After isolating homozygous individuals, *ipt3* knockout was confirmed by qRT-PCR analyses (**Fig. 3B**). Then, *ipt3* knockout mutants and wild-type plants were used to perform nodulation assays. At 14 dai, all the *ipt3* knockout mutants showed a significantly lower nodule number than the control (**Fig. 3C and D**). Moreover, between 40-55% of the *ipt3* mutants showed neither emerging nodule primordia nor nodules, while only 15% of wild-type plants lacked nodules (**Fig. 3E**). Root dry weight measurements revealed no significant differences between wild-type plants and *ipt3* mutants after 14 dai (**Fig. 3F**), while shoot dry weight significantly decreased only in *ipt3-1* (**Fig. 3G**). These results strongly suggest that the lower nodule number observed in *ipt3* mutants is a consequence of the loss of IPT3 function and not a pleiotropic effect caused by *Tnt1* insertions.

**Figure 3.**
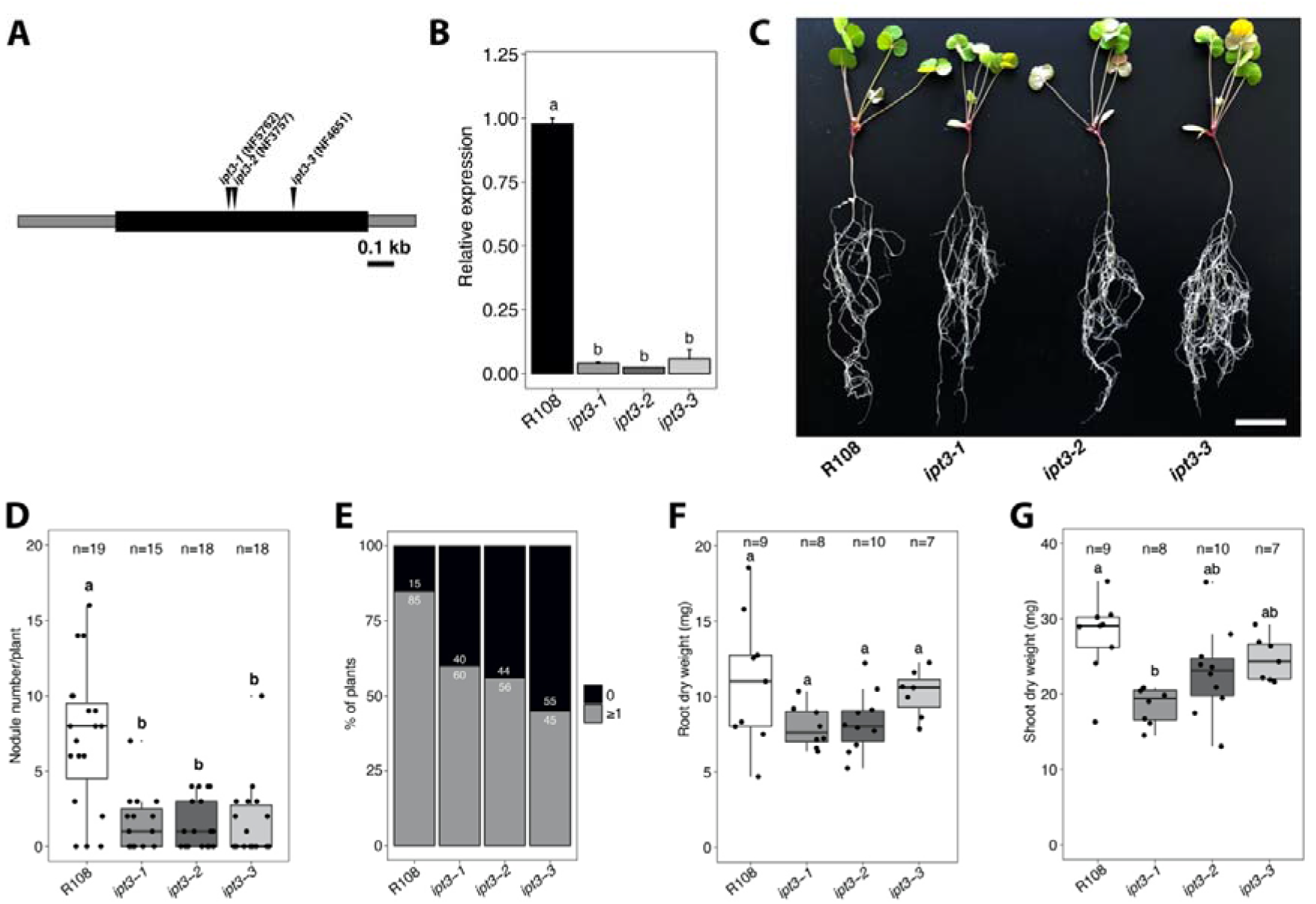
*IPT3* is required for nodule development in *M. truncatula*. (A) Schematic diagram showing genomic *IPT3* gene structure. NF5762, NF3757, and NF4651 *Tnt1* lines have one insertion in the unique exon region (black) flanked by 5’and 3’ untranslated regions (gray) of the gene and were renamed *ipt3-1, ipt3-2*, and *ipt3-3*, respectively. Black bar, 100 bp. (B) qRT-PCR analysis of *IPT3* gene expression in R108 genotype and *ipt3* homozygous mutants. Values indicates means ± SE for three biological replicates (n=3). *P* values were calculated by ANOVA followed by Tukey’s posthoc testing. Groups of different significance, at least *P*<0.05, are indicated with different letters. (C) Representative image of 3-week-old R108 and *ipt3* mutant plants after 14 dai of treatment with *S. meliloti* 1021. White bar represents 3 cm. (D) Nodule number in R108 and *ipt3* mutants after 14 dai. Statistical analysis was performed using ANOVA followed by Tukey’s posthoc testing. Groups of different significance, at least *P*<0.05, are indicated with different letters. (E) Percentage of wild-type and *ipt3* mutant plants showing 1 or more nodules or none after 14 dai. (F) Root and (G) shoot dry weight measurements of wild-type and *ipt3* mutant plants after 14 dai. Statistical analysis was performed using ANOVA followed by Tukey’s posthoc testing. Groups of different significance, at least *P*<0.05, are indicated with different letters.

### *IPT3* expression is induced in the stele at the base of the developing nodule primordium

To get further insights into the role of *IPT3* during nodule organogenesis, we investigated *IPT3* expression by monitoring the activity of tdTOMATO fluorescent protein driven by *IPT3* promoter in a time-course experiment. This strategy allowed us to monitor *IPT3* expression and CK signaling simultaneously after the inoculation with *S. meliloti* through live imaging in *M. truncatula* transgenic roots. To achieve this goal, we cloned the *IPT3* promoter (2312 bp; see Method section) in frame with the tdTOMATO fluorescent protein fused to a nuclear localization signal peptide resulting in the *pIPT3::nls:tdTOMATO* construction. Before inoculation with rhizobia, *IPT3* expression and CK response overlapped at the root stele (**Fig. S3 A-C**). At 24 hours post *S. meliloti* inoculation, *IPT3* expression was similar to the control treatment and still localized in the vasculature (**Fig. S3 D-F**). In contrast, the CK response was observed in the epidermis and outer cortical cells of the SZ, as described above (**Fig. S3 G-I**). At 2 dai, CK response expanded to the majority of root cortical layers of the SZ, while *pIPT3::nls:tdTOMATO* activity was still mainly found in the stele with a similar expression level to that of the control (**Fig. 4A-C; Fig S3E and H**). This suggests that IPT3 is not involved in the CK signaling activation during the early symbiotic interaction described in the present research work (48 hai).

**Figure 4.**
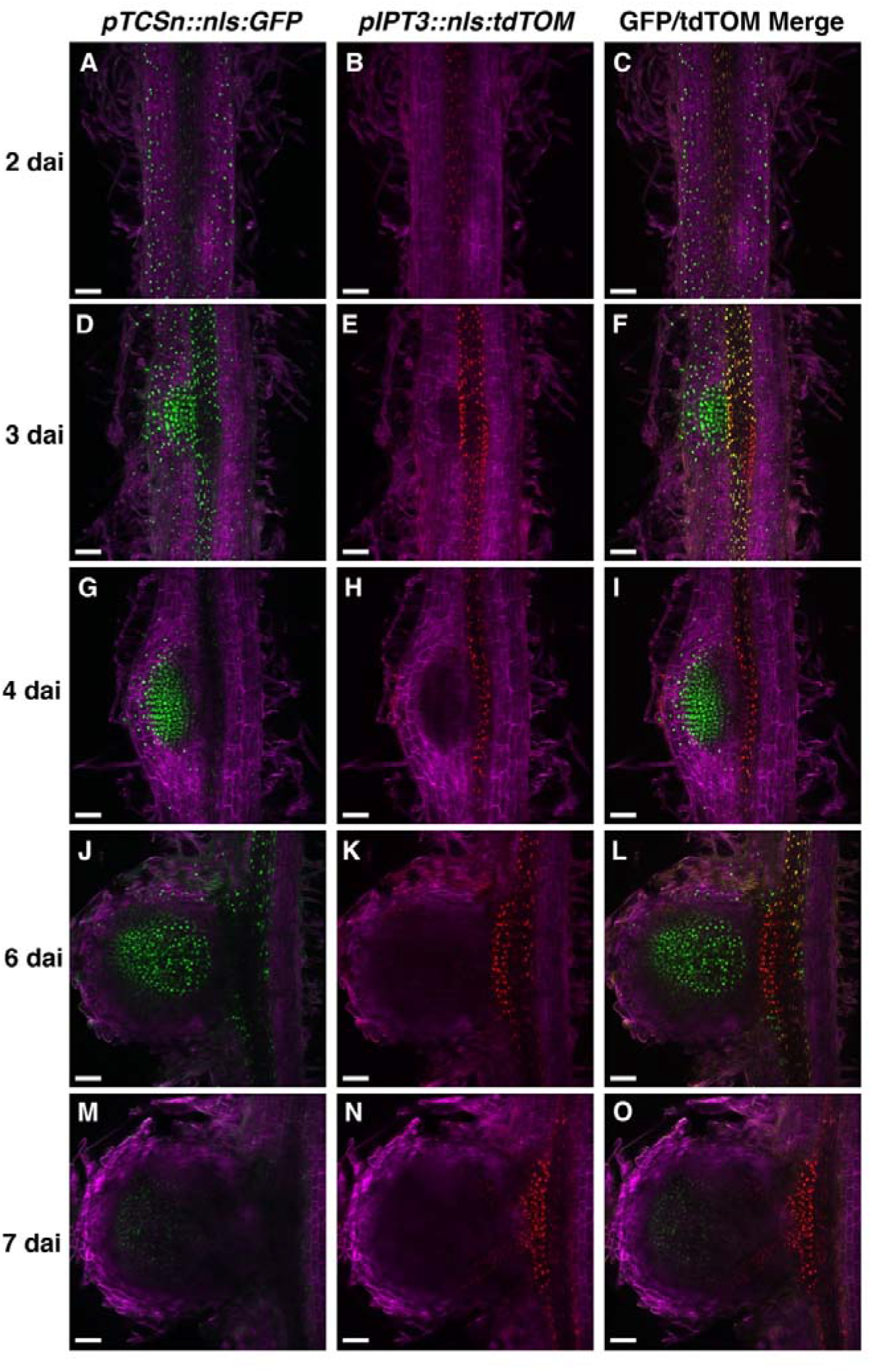
*IPT3* expression is induced in the stele at the base of the nodule primordium during the first cortical cell divisions. (A-C)*pTCSn::nls:tGFP* and *pIPT3::nls:tdTOMATO* activities in the susceptible zone of transgenic root after 2 dai of *S. meliloti*. (D-I) *pTCSn::nls:tGFP* and *pIPT3::nls:tdTOMATO* activities in different developmental stages of nodule primordium after 3 (D-F) and 4 dai (G-I). (J-O) *pTCSn::nls:tGFP* and *pIPT3::nls:tdTOMATO* activities in different developmental stages of mature nodules after 6 (J-L) and 7 dai (M-O). Green and red represent fluorescence signals emitted by tGFP and tdTOMATO, respectively. The white bar shows 100 μm.

At 3 dai, nodule primordium was initiated (between stage II and III), and CK response was mainly localized to the dividing cortical cells and stele **(Fig 4D)**. *IPT3* expression was strongly induced in the stele at the base of dividing cortical cells of the nodule primordium **(Fig. 4E)**. This result is consistent with prior reports of the induction of the *IPT3* transcription 72 hours after *S. meliloti* inoculation (Schiessl et al., 2019; **Fig. S4**). The induction of the CK response and *IPT3* expression overlapped at the root vasculature and the base of the nodule primordium, likely involving pericycle **(Fig. 4F)**. At 4 dai, with the nodule primordium at advanced stages of development and CK response extended to more cortical cell layers **(Fig. 4G)**, *IPT3* expression levels remained high and localized in the stele below the dividing cortical cells **(Fig. 4H)**, showing lower overlapping with CK signaling in the stele **(Fig. 4I)**. At 6 dai, CK response was mainly localized at the central zone of the nodule **(Fig. 4J)**, while *IPT3* expression was strongly activated and started to propagate from the root stele to the nodule vasculature **(Fig. 4K)**. At this stage, the nodule meristem activates, and the overlap between the CK response and *IPT3* expression declined, with each one showing a specific spatial pattern **(Fig. 4L)**. At 7 dai, CK response was mainly localized to the nodule meristem **(Fig 4M)**. At the same time, *IPT3* expression was also detected in the developing nodule vasculature **(Fig. 4N)**, in line with *IPT3* expression in nodule vascular bundles recently reported using *pIPT3::GUS* (Azarakhsh et al., 2020). At this stage, CK response was restricted to the nodule meristem, and *IPT3* expression was localized at the base of the nodule and vascular bundles **(Fig. 4O)**. Together, these results suggest that the *IPT3* induction at the base of the nodule primordium contributes to the biosynthesis of CK, which in turn triggers CK signaling during nodule organogenesis.

### *IPT3* is required for the induction of symbiotic genes during nodule initiation

Rhizobia-induced nodule initiation is dependent on crucial nodule development regulators that promote cortical cell division, including the cytokinin receptor CRE1, the transcription factor NIN and its targets LBD16 and NFYA1 (Gonzalez-Rizzo et al., 2006; Plet et al., 2011; Laporte et al., 2014; Vernié et al., 2015; Schiessl et al., 2019). *ipt3* mutants show impairment of nodule development **(Fig. 3D)**, suggesting that the biosynthesis of CK precursors by IPT3 is required to activate CK-induced positive regulators of nodule development. To test this hypothesis, we performed an *in vitro* nodulation assay using wild-type and two *ipt3* mutant lines, *ipt3-2* and *ipt3-3*, to compare gene expression of these essential regulatory genes. *ipt3-1* was excluded due to the previously mentioned reduced shoot mass phenotype. 3-day-old *M. truncatula* plants were inoculated with *S. meliloti*, and the SZ of the root was harvested at 4 dai for gene expression analyses. We found that expression of *NIN*, *LBD16*, and *NFYA1* was upregulated in wild-type compared to non-inoculated control plants. In contrast, in the *ipt3* mutants, these genes were not significantly induced **(Fig. 5)**, indicating that the transcriptional activation of positive regulators of nodulation requires IPT3 at 4 dai. *CYCLINA3;A* (*CYC3;A*), a CK-induced gene and a cell division marker during nodule initiation (Schiessl et al., 2019), did not show significant induction either in the wild-type or *ipt3* mutants with respect to the control treatment **(Fig. 5)**. Besides *CRE1*, we found significant transcriptional induction of CK signaling genes, such as *RRA3* and *RRA11*, which was affected in *ipt3* mutants compared to the control **(Fig. 5)**, indicating IPT3 is required for the activation of CK signaling during nodule initiation. On the contrary, *RRA4* was not induced in wild-type and *ipt3* mutants by rhizobia compared to the control **(Fig. 5)**. *IPT3* was also upregulated in wild-type plants at 4 dai with respect to the control **(Fig. 5)**, consistent with the rhizobia-induced expression pattern observed in previous work and with the visual reporter (**Fig. S4**; Schiessl et al., 2019; **Fig. 4H**). Together, these results suggested that rhizobia-induced *IPT3* expression contributes to CK biosynthesis. In turn, CK biosynthesis promotes transcriptional activation of positive nodule development regulators and CK signaling genes required for nodule development.

**Figure 5.**
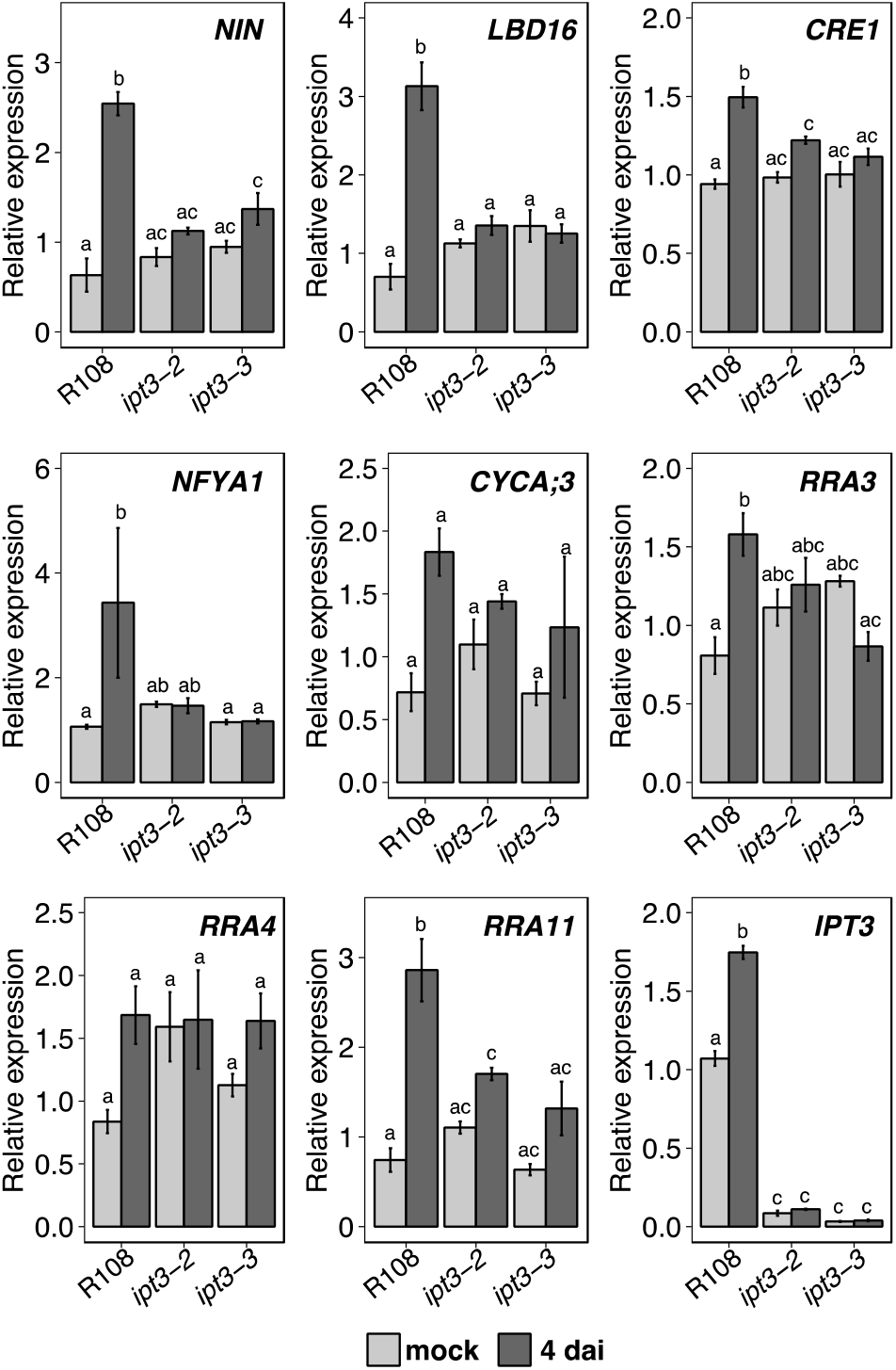
Rhizobia-dependent induction of nodulation regulators and CK signaling genes is impaired in *ipt3* loss-of-function mutants. qRT-PCR analyses of nodulation regulators (*NIN, LBD16, NFYA1*) cell cycle (*CYCA;3*) and CK signaling genes (*CRE1, RRA3, RRA4, RRA11*) after mock treatment or 4 dai in (A) wild-type, (B) *ipt3-2* and (C) *ipt3-3* mutants. Values are the mean of fold-changes of three biological replicates normalized to the untreated wild-type value (set as 1) for each gene for all genotypes. Values indicate means ± SE of three biological replicates (n=3). *P* values were calculated by ANOVA followed by Tukey’s posthoc testing. Groups of different significance, at least *P*<0.05, are indicated with different letters.

## DISCUSSION

In the present study we developed a novel tGFP-based *TCSn* reporter that allowed us to perform live imaging tissue-specific time course, with a high temporal resolution of the rhizobia-induced CK signaling. This reporter system allowed us to gain further insights into the cytokinin signaling induction during rhizobia perception and nodule formation in *M. truncatula*. Our data indicate that the CK signaling activation occurs in multiple, discrete stages, initially activating in the epidermis of the root SZ and expanding across the cortex during the first 48 hrs. Rhizobia trigger this first wave of CK signaling in the SZ of the *M. truncatula* root. After 48 hours, this widespread CK signaling activation disappears, giving way to the second wave of CK signaling activation in the cortex, localized in the specific zones where the cell divisions that will give rise to the nodule primordium. The characterization of *ipt3* mutants suggests that IPT3 contributes to the CK biosynthesis that triggers this second wave of signaling activation.

It has been proposed recently that the *MtABCG56* transporter, which is transcriptionally induced between 6 and 24 hours after the LCO treatment, exports bioactive CKs from the root epidermis to the cortex, promoting CRE1-dependent cortical CK response (Jarzyniak et al., 2021). In agreement with this model, we observed that CK signaling was activated at 24 hai in the outer cortical cells **(Fig. 2C)**, and it extends to the majority of cortical cell layers at 48 hai in the SZ **(Fig. 2D)**, possibly due to the amplification of CRE1-triggered CK response. These results are consistent with the similar CK activation patterns observed previously using *pTCSn:GUS*, showing GUS activity localized at the epidermis and outer cortical cells at 8 hai and the inner cortical cells at 72 hai, and *in situ* hybridization detected expression of *RRA4* mRNA widely in the root cortex at 48 hai (Vernié et al., 2015; Jardinaud et al., 2016). Here, a live imaging time course allowed us to precisely elucidate the timing of the CK signaling activation pattern observed during the early symbiosis interaction. Consistent with these observations, *in situ* hybridization showed that *RR4* mRNA levels strongly accumulate in the different *M. truncatula* root cortex layers at 48 hai with *S. meliloti* (Vernié et al., 2015). In contrast, in *L. japonicus*, CK signaling activation in cortical cells precedes epidermal CK responses (Held et al., 2014; Reid et al., 2017). These results suggest that the spatial-temporal CK signaling activation may differ between determinate and indeterminate nodulating species. The difference in rhizobia-triggered CK signaling patterns between these species highlights the need for high-resolution tissue-specific characterization of the CK responses in other legumes.

The second wave of CK signaling activation in the cortex requires *de novo* CK biosynthesis. It has been reported that CK biosynthesis genes, such as *LOG1, LOG2*, and *IPT3*, are expressed in the nodule primordium of *M. truncatula* (Mortier et al., 2014; Azarakhsh et al., 2020). It has been proposed that KNOX3 transcription factor directly promotes the expression of cytokinin biosynthesis genes by *MtLOG1, MtLOG2*, and *MtIPT3* which are co-expressed during nodule development in *M. truncatula* (Azarakhsh et al., 2020). By monitoring the *pIPT3::nls:tdTOMATO* and the CK sensor simultaneously, we resolved the spatial-temporal pattern of *IPT3* expression and its interaction with the CK signaling during the indeterminate nodule development in *M. truncatula*. We found that *IPT3* is induced in the stele, likely at the pericycle and adjacent cells to the first dividing cortical cells at 3 dai **(Fig. 4E)**, overlapping with CK signaling activation at the base of the nodule primordium **(Fig. 4F)**. This result disagrees with the *pIPT3::GUS* activity reported in the whole nodule primordium after 3 days post inoculation (Azarakhsh et al., 2020). This discrepancy may be explained by a higher sensitivity of the GUS reporter, which could reveal the promoter activity even with very low expression levels. This outcome may also be derived by the diffusion of the GUS reaction product to adjacent cells, especially when extended incubation periods are required. However, similarly to what was reported with *pIPT3::GUS*, after 7 days post inoculation (Azarakhsh et al., 2020), we found that *IPT3* expression spreads from the stele at the base of the nodule to the nodule developing vascular bundles **(Fig. 4N)**.

IPT enzymes catalyze the formation of iP riboside 5’-diphosphate (iPRDP) or iP riboside 5’-triphosphate (iPRTP), which are precursors for bioactive cytokinin biosynthesis by LOG family enzymes (Sakakibara, 2006). The induction of *IPT3* at the base of the nodule primordium may provide the substrate for *LOG* genes, such as *LOG1* and *LOG2*, which catalyzes the production of the bioactive CKs (Kurakawa et al., 2007). Indeed, analyses of *LOG1* and *LOG2* promoters fused to GUS revealed that these genes are expressed in the central zone of developing nodule primordia, overlapping with our observations of the CK signaling activation (Mortier et al., 2014; Azarakhsh et al., 2020). Similarly, *LOG1* RNAi plants show lower nodules than control plants in *M. truncatula* (Mortier et al., 2014), and we found a similar result in our *ipt3* loss of function mutants. The same effect has been reported in *L. japonicus IPT3* RNAi plants, which produced fewer ITs and nodules than wild-type (Chen et al., 2014). However, it has been proposed that *LjIPT3* also participates in the generation of shoot-derived CK precursors, which participate in the autoinhibition of nodulation (AON) mechanism in *L. japonicus* (Sasaki et al., 2014). These findings suggested that IPT3-derived CKs may be involved in different mechanisms in shoots and roots. The different timing of *IPT3* activation explains its dual role during nodulation of *L. japonicus*. If *IPT3* participates in AON in *M. truncatula* remains an open question.

Rhizobia-dependent transcriptional activation of CK-induced positive regulators of nodule development, such as *NIN, CRE1, LBD16*, and *NFYA1*, was affected in *ipt3* mutants at 4 dai **(Fig. 5)**. Together, these results suggested that *de novo* CK precursor synthesis is required for the CK-mediated induction of key nodule development regulators, promoting CK downstream responses and cell divisions during nodule development.

In conclusion, we propose a high-resolution model for the tissue-specific temporal activation pattern of CK signaling during indeterminate nodule development in *M. truncatula* **(Fig. 6)**. Furthermore, we show that *IPT3*, a cytokinin biosynthesis gene, contributes to *de novo* CK biosynthesis, which in turn promotes the expression of a critical positive regulator of nodule development, such as *NIN* and *CRE1*.

**Figure 6.**
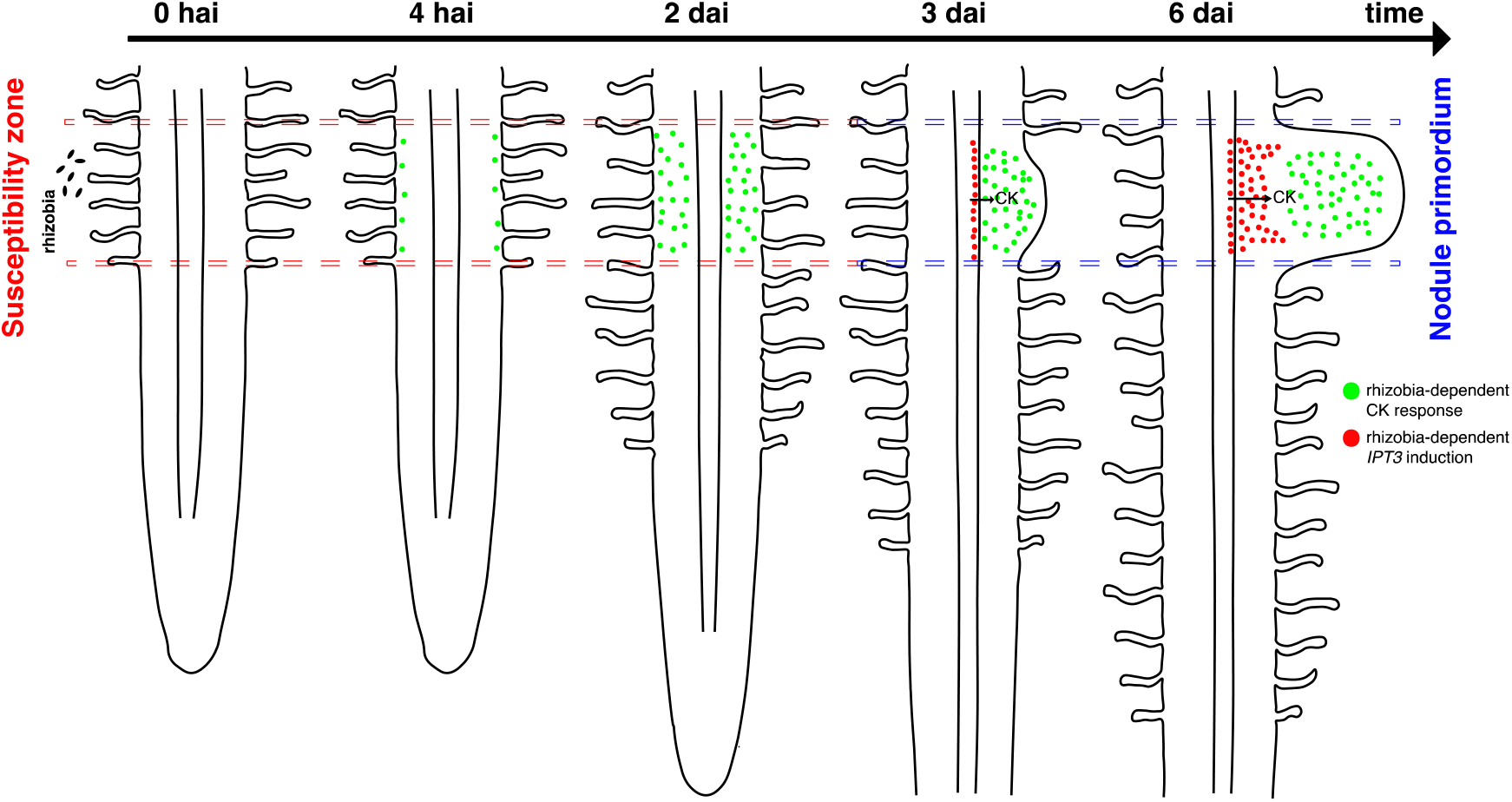
Schematic representation of the spatiotemporal regulation of CK response induced by rhizobia and the proposed function of IPT3 during indeterminate nodule development in *M. truncatula*. At 4 hai, CK signaling activation starts in epidermal cells, and progress to the majority of cortical cell layers within 48 hrs after rhizobia perception. At 3 dai, CK signaling is activated and localized in dividing cortical cells and *IPT3* expression is induced in the stele at the base of nodule primordium. At 6 dai, CK response is localized in the central zone of nodule primordium and *IPT3* is strongly activated and started to propagate from the root stele to the nodule vasculature.

## MATERIALS AND METHODS

### Plant material and growth conditions

*Medicago truncatula* R108 is the wild-type background for the *Tnt1* mutant lines; Jemalong A17 was used for the rest. Seeds were scarified 8 min in sulfuric acid and sterilized for 4 min in bleach (12% [v/v] sodium hypochlorite). After rinsing with sterilized water, seeds were sown on 1% agar plates supplemented with 1 μM gibberellic acid and stored at 4°C for 3 days before incubating overnight at 24°C in the dark. Germinated seedlings were transferred to square plates (22.5×22.5 cm) containing modified Fahräeus medium (Boisson-Dernier et al., 2001) supplemented with 15 mM NH4NO3 and grown vertically at 24°C under long-day conditions in a growth chamber (16 h light/8 h dark; 150 μmol m^−2^ s^−1^ light intensity).

### Cloning

The Golden Gate MoClo and MoClo Plant toolkits (New England Biolabs) were used to obtain all constructions described in this work (Engler et al., 2014). The cloning reaction consists of 25 cycles of 3 min at 37°C for digestion and 4 min at 16°C for ligation combining the restriction enzymes *Bsa*I-HF (New England BioLabs, Ipswich, UK) for level 1 reaction or *Bpi*I (Thermo Fisher Scientific, Waltham, USA) for level 2 reaction together with T4 DNA Ligase (New England BioLabs, Ipswich, UK). *TCSn* promoter DNA sequence (Zürcher et al., 2013) comprising overhangs was synthesized by Synbio Technologies and cloned in frame with a nuclear localization signal (Addgene, catalog number: 50294), turbo GFP (Addgene, catalog number: 50322), and *Solanum lycopersicum* ATPase terminator (Addgene, catalog number: 50344) in the level 1 vector pICH47811 (Addgene, catalog number: 48008). To amplify *IPT3* promoter (gene ID v4: Medtr1g072540; v5: MtrunA17Chr1g0185751) genomic DNA (gDNA) extraction was performed as previously described (Causevic et al., 2005) with the following modifications. 3-day-old *M. truncatula* A17 seedlings were frozen in liquid nitrogen and ground in 2 ml tubes using a homogenizer (MiniG 1600, Spex, Metuchen, NJ). 200 μL of pre-heated CTAB buffer containing 0.2% β-mercaptoethanol, 4% PVP, and 0.1 mg/ml Proteinase K (Invitrogen) was added to the ground tissue and digested for 1 h at 60 °C. After adding 1 volume of chloroform/isoamyl alcohol (24:1, v/v), samples were mixed and then centrifuged at 14,000xg for 10 min. The aqueous phase was transferred to a new tube and incubated with 1 μL of RNase A (Macherey-Nagel, 12mg/mL stock) for 30 min at 37 °C. The chloroform/isoamyl alcohol cleaning step was repeated and the aqueous phase was transferred to a new tube and mixed with 1 volume of isopropanol and incubated for 30 min at −20 °C. The sample was centrifuged at 14,000xg for 15 min, and the pellet was washed with 70% and 95% ethanol by centrifugation for 5 min. The pellet was air-dried and resuspended in 50 μL of TE buffer (Invitrogen, catalog number: 12090015). A genomic fragment of 2312 bp upstream of the *IPT3* coding sequence including the 5’ UTR was amplified using primers containing Golden Gate overhangs **(Table S1)** from *M. truncatula* A17 gDNA. 100 ng of gDNA were used to set up a PCR using Phusion^®^ High-Fidelity DNA Polymerase (New England BioLabs). The PCR product was separated in a 1% agarose gel by electrophoresis. *IPT3* promoter DNA sequence was extracted and purified from the gel using Monarch DNA Gel Extraction Kit (New England Biolabs) and Sanger sequenced at Genewiz (South Plainfield, NJ). Also, tdTOMATO coding sequence (CDS) was amplified using primers containing Golden Gate overhangs **(Table S1)**. IPT3 promoter was cloned in frame with a nuclear localization signal (Addgene, catalog number: 50294), tdTOMATO CDS, and 35S terminator (Addgene, catalog number: 50337) in the level 1 vector pICH47802 (Addgene, catalog number: 48007). Level 1 *pICH47811-pTCSn::nls:tGFP::tATPase* (position 2) was cloned together with *pICH47802-p35S::ER:tdTOM::tNOS* selection marker (position 1) in the final plant expression vector pAGM4673 (Addgene, catalog number: 48014). For simultaneous live imaging, *pICH47811-pTCSn::nls:tGFP* (position 2) was cloned together with *pICH47802-pIPT3::nls:tdTOMATO::t35S* (position 1) in the final plant expression vector pAGM4673 (Addgene, catalog number: 48014).

### Genotyping of *Tnt1* insertion lines

*M. truncatula* R108 *Tnt1* transposon insertion lines utilized in this research project, which are jointly owned by the Centre National De La Recherche Scientifique, were obtained from the Noble Research Institute, and were created through research funded, in part, by grants from the National Science Foundation, NSF #DBI-0703285, and NSF #IOS-1127155. Three different *Tnt1* transposon insertion lines, namely NF5762 (*ipt3-1*), NF3757 (*ipt3-2*), and NF4651 (*ipt3-3*), were genotyped by PCR using *Tnt1*-specific primers (Cheng et al., 2011) combined with *MtIPT3* gene-specific primers encompassing the insertions **(Table S1)**. The expression of *MtIPT3* in homozygous plants was tested by RT–qPCR to confirm they were knockout mutants for the *MtIPT3* gene **(Table S1)**.

### *Agrobacterium rhizogenes*-mediated transformation and *in-vitro* nodulation assay

The constructs described above were introduced into *Agrobacterium rhizogenes* MSU440 electrocompetent cells and used to generate transgenic roots in *M. truncatula* (Boisson-Dernier et al., 2001). The transgenic roots were selected based on the fluorescence emitted by *pTCSn::nls:tGFP* construct at the root tip under a fluorescence stereomicroscope. For *in-vitro* nodulation assay time-course experiment, 4-week-old transgenic roots were transferred to Buffered Nodulation Medium (BNM) (Ehrhardt et al., 1992) supplemented with 0.1 μM AVG (aminoethoxyvinyl glycine hydrochloride; Sigma-Aldrich) to reduce ethylene production. After 5 days of acclimation, transgenic roots were treated with a suspension of *S. meliloti* (OD_600_=0.02) supplemented with 3 μM of luteolin to activate LCO production (Sigma-Aldrich) collecting transgenic roots at different timepoints.

### *In vitro* Cytokinin treatment and *S. meliloti* inoculation

3-day-old *M. truncatula* seedlings were transferred to Fahräeus medium supplemented with water (mock treatment) or 1 μM of 6-BAP (6-benzylaminopurine; Sigma-Aldrich) and maintained under the same growth conditions. Five roots from different seedlings were collected after 1, 8, 24, and 48 hours after CK treatment and immediately frozen in liquid nitrogen for RNA extraction. To analyze *pTCSn::nls:tGFP* activity after CK treatment, *M. truncatula* transgenic roots were submerged with a solution of 1 μM 6-BAP for 5 min and then the solution was poured off. After 24 hours, transgenic roots harboring *pTCSn::nls:tGFP* construct were harvested and used for microscopic analysis. For qRT-PCR studies of nodule development regulator and CK signaling genes in wild-type and *ipt3* mutants, germinated seedlings were transferred to nitrogen-free Fahräeus medium and grown under the same conditions described above. 3-day-old seedlings were inoculated alongside the root with 200 μL/root of *S. meliloti* (OD_600_=0.02) resuspended in liquid nitrogen-free modified Fahräeus medium or with Fahräeus medium (mock treatment). Four days after inoculation, root segments from the susceptibility zone were harvested and pooled (10 plants per sample) for RNA extraction.

### Gene expression analysis

RNA extraction was performed using as previously described (Chang et al., 1993). RNA samples were digested with DNase I and purified using RNA Clean & Concentrator™-5 kit (Zymo Research, Irvine, CA). Quantitative reverse transcription-polymerase chain reaction (qRT-PCR) analyses were performed using Luna Universal Probe One-Step qRT-PCR Kit (New England Biolabs) following the manufacturer’s instructions, using 200 ng of total RNA. *EF1α* gene was used as a housekeeping gene and the average of two technical replicates was obtained to calculate relative gene expression (ΔΔCt method). Primers used for qRT-PCR experiments are listed in **Table S1**.

### Microscopic live imaging

Before microscopic imaging, transgenic roots were placed in a staining solution with 0.1% Calcofluor white M2R (Sigma-Aldrich) in PBS 1X (Corning) for 10 minutes at room temperature and then rinsed with PBS 1X before imaging. A confocal microscope Leica TCS SP5 confocal 119 was used for live imaging. At least 10 different roots for treatment or time point were analyzed. The excitation wavelengths for tGFP, tdTOMATO, and Calcofluor white M2R were 488, 561, and 405 nm, respectively. Fluorescent signals were collected at 497–557nm (tGFP), 570–643nm (tdTOMATO), or 450–480 nm (Calcofluor white M2R). Fluorescent signals are presented with green (tGFP), red (tdTOMATO), and magenta (Calcofluor white M2R). For DNA staining, transgenic roots were incubated with 5 μg/ml 4’,6-diamidino-2-phenylindole (DAPI) in PBS 1X (Corning) for 10 min at room temperature and then rinsed with PBS 1X before imaging. DAPI excitation wavelength was 405 nm, while emission was collected at 450–480 nm.

### *In vivo* Nodulation assay and dry weight measurement

For *in vivo* nodulation assay, seedlings were germinated as described above. Then, plants were grown in pots (9×9×9 cm) containing pre-sterilized calcined clay, Turface^®^ (Profile Products, Buffalo Grove, IL) and sand (2:2 v/v), where Turface^®^ was placed at the bottom and on the top of a layer of sand. Plants were watered with modified Fahräeus medium supplemented with 0.5 mM of NH4NO3 and covered with a lid. After one week of acclimation, Fahräeus medium supplemented with 0.5 mM of NH_4_NO_3_ was removed entirely from the tray and replaced with a nitrogen-free modified Fahräeus medium. Plants were treated pouring 10 mL of *S. meliloti* 1021 suspension (OD_600_=0.02) into each pot. Plants were watered using nitrogen-free modified Fahräeus medium every 2-3 days. After 2 weeks of inoculation, nodule number was assessed inspecting plant roots under a stereomicroscope. After counting nodules, roots and shoots for each individual were separated and dried in paper bags. After 48 hours of drying at 65°C, root and shoot dry weight were recorded.

## Parsed Citations

Ariel F, Brault-Hernandez M, Laffont C, Huault E, Brault M, Plet J, Moison M, Blanchet S, Ichanté JL, Chabaud M, et al (2012) Two Direct Targets of Cytokinin Signaling Regulate Symbiotic Nodulation in Medicago truncatula. Plant Cell 24: 3838–3852

Azarakhsh M, Kirienko AN, Zhukov VA, Lebedeva MA, Dolgikh EA, Lutova LA (2015) KNOTTED1-LIKE HOMEOBOX 3: a new regulator of symbiotic nodule development. EXBOTJ 66: 7181–7195

Azarakhsh M, Lebedeva MA, Lutova LA (2018) Identification and Expression Analysis of Medicago truncatula Isopentenyl Transferase Genes (IPTs) Involved in Local and Systemic Control of Nodulation. Front Plant Sci 9: 304

Azarakhsh M, Rumyantsev AM, Lebedeva MA, Lutova LA (2020) Cytokinin biosynthesis genes expressed during nodule organogenesis are directly regulated by the KNOX3 protein in Medicago truncatula. PLoS ONE 15: e0232352

Boisson-Dernier A, Chabaud M, Garcia F, Bécard G, Rosenberg C, Barker DG (2001) Agrobacterium rhizogenes-Transformed Roots of Medicago truncatula for the Study of Nitrogen-Fixing and Endomycorrhizal Symbiotic Associations. MPMI 14: 695–700

Boivin S, Kazmierczak T, Brault M, Wen J, Gamas P, Mysore KS, Frugier F (2016) Different cytokinin histidine kinase receptors regulate nodule initiation as well as later nodule developmental stages in Medicago truncatula: Specificity and redundancy of CHKs in nodulation. Plant, Cell & Environment 39: 2198–2209

Buhian WP, Bensmihen S (2018) Mini-Review: Nod Factor Regulation of Phytohormone Signaling and Homeostasis During Rhizobia-Legume Symbiosis. Front Plant Sci 9: 1247

Causevic A, Delaunay A, Ounnar S, Righezza M, Delmotte F, Brignolas F, Hagège D, Maury S (2005) DNA methylating and demethylating treatments modify phenotype and cell wall differentiation state in sugarbeet cell lines. Plant Physiology and Biochemistry 43: 681–691

Chang S, Puryear J, Cairney J (1993) A simple and efficient method for isolating RNA from pine trees. Plant Mol Biol Rep 11: 113–116

Chen Y, Chen W, Li X, Jiang H, Wu P, Xia K, Yang Y, Wu G (2014) Knockdown of LjIPT3 influences nodule development in Lotus japonicus. Plant and Cell Physiology 55: 183–193

Cheng X, Wen J, Tadege M, Ratet P, Mysore KS (2011) Reverse Genetics in Medicago truncatula Using Tnt1 Insertion Mutants. Methods in Molecular Biology 678:179–90.

Damiani I, Drain A, Guichard M, Balzergue S, Boscari A, Boyer J-C, Brunaud V, Cottaz S, Rancurel C, Da Rocha M, et al (2016) Nod Factor Effects on Root Hair-Specific Transcriptome of Medicago truncatula: Focus on Plasma Membrane Transport Systems and Reactive Oxygen Species Networks. Front Plant Sci. doi: 10.3389/fpls.2016.00794

Ehrhardt D, Atkinson E, Long (1992) Depolarization of alfalfa root hair membrane potential by Rhizobium meliloti Nod factors. Science 256: 998–1000

Engler C, Youles M, Gruetzner R, Ehnert T-M, Werner S, Jones JDG, Patron NJ, Marillonnet S (2014) A Golden Gate Modular Cloning Toolbox for Plants. ACS Synth Biol 3: 839–843

Ferguson BJ, Indrasumunar A, Hayashi S, Lin M-H, Lin Y-H, Reid DE, Gresshoff PM (2010) Molecular Analysis of Legume Nodule Development and Autoregulation. Journal of Integrative Plant Biology 52: 61–76

Ferguson BJ, Mathesius U (2014) Phytohormone Regulation of Legume-Rhizobia Interactions. J Chem Ecol 40: 770–790

Fisher J, Gaillard P, Fellbaum CR, Subramanian S, Smith S (2018) Quantitative 3D imaging of cell level auxin and cytokinin response ratios in soybean roots and nodules: quantitative imaging of auxin and cytokinin. Plant Cell Environ. doi: 10.1111/pce.13169

Fonouni-Farde C, Kisiala A, Brault M, Emery RJN, Diet A, Frugier F (2017) DELLA1-Mediated Gibberellin Signaling Regulates Cytokinin-Dependent Symbiotic Nodulation. Plant Physiol 175: 1795–1806

Gamas P, Brault M, Jardinaud M-F, Frugier F (2017) Cytokinins in Symbiotic Nodulation: When, Where, What For? Trends in Plant Science 22: 792–802

Gonzalez-Rizzo S, Crespi M, Frugier F (2006) The Medicago truncatula CRE1 Cytokinin Receptor Regulates Lateral Root Development and Early Symbiotic Interaction with Sinorhizobium meliloti. Plant Cell 18: 2680–2693

Heckmann AB, Sandal N, Bek AS, Madsen LH, Jurkiewicz A, Nielsen MW, Tirichine L, Stougaard J (2011) Cytokinin Induction of Root Nodule Primordia in Lotus japonicus Is Regulated by a Mechanism Operating in the Root Cortex. MPMI 24: 1385–1395

Held M, Hou H, Miri M, Huynh C, Ross L, Hossain MS, Sato S, Tabata S, Perry J, Wang TL, et al (2014) Lotus japonicus Cytokinin Receptors Work Partially Redundantly to Mediate Nodule Formation. Plant Cell 26: 678–694

Hossain MS, Shrestha A, Zhong S, Miri M, Austin RS, Sato S, Ross L, Huebert T, Tromas A, Torres-Jerez I, et al (2016) Lotus japonicus NF-YA1 Plays an Essential Role During Nodule Differentiation and Targets Members of the SHI/STY Gene Family. MPMI 29: 950–964

Jardinaud M-F, Boivin S, Rodde N, Catrice O, Kisiala A, Lepage A, Moreau S, Roux B, Cottret L, Sallet E, et al (2016) A Laser Dissection-RNAseq Analysis Highlights the Activation of Cytokinin Pathways by Nod Factors in the Medicago truncatula Root Epidermis. Plant Physiol 171: 2256–2276

Jarzyniak K, Banasiak J, Jamruszka T, Pawela A, Di Donato M, Novák O, Geisler M, Jasiński M (2021) Early stages of legume–rhizobia symbiosis are controlled by ABCG-mediated transport of active cytokinins. Nat Plants. doi: 10.1038/s41477-021-00873-6

Kieber JJ, Schaller GE (2018) Cytokinin signaling in plant development. Development 145: dev149344

Kurakawa T, Ueda N, Maekawa M, Kobayashi K, Kojima M, Nagato Y, Sakakibara H, Kyozuka J (2007) Direct control of shoot meristem activity by a cytokinin-activating enzyme. Nature 445: 652–655

Laloum T, Baudin M, Frances L, Lepage A, Billault-Penneteau B, Cerri MR, Ariel F, Jardinaud M-F, Gamas P, de Carvalho-Niebel F, et al (2014) Two CCAAT-box-binding transcription factors redundantly regulate early steps of the legume-rhizobia endosymbiosis. Plant J 79: 757–768

Laporte P, Lepage A, Fournier J, Catrice O, Moreau S, Jardinaud M-F, Mun J-H, Larrainzar E, Cook DR, Gamas P, et al (2014) The CCAAT box-binding transcription factor NF-YA1 controls rhizobial infection. EXBOTJ 65: 481–494

Liu C-W, Breakspear A, Roy S, Murray JD (2015) Cytokinin responses counterpoint auxin signaling during rhizobial infection. Plant Signaling & Behavior 10: e1019982

Madsen LH, Tirichine L, Jurkiewicz A, Sullivan JT, Heckmann AB, Bek AS, Ronson CW, James EK, Stougaard J (2010) The molecular network governing nodule organogenesis and infection in the model legume Lotus japonicus. Nat Commun 1: 10

Mortier V, Wasson A, Jaworek P, De Keyser A, Decroos M, Holsters M, Tarkowski P, Mathesius U, Goormachtig S (2014) Role of LONELY GUY genes in indeterminate nodulation on Medicago truncatula. New Phytol 202: 582–593

Op den Camp RHM, De Mita S, Lillo A, Cao Q, Limpens E, Bisseling T, Geurts R (2011) A Phylogenetic Strategy Based on a Legume-Specific Whole Genome Duplication Yields Symbiotic Cytokinin Type-A Response Regulators. Plant Physiol 157: 2013–2022

Ovchinnikova E, Journet E-P, Chabaud M, Cosson V, Ratet P, Duc G, Fedorova E, Liu W, den Camp RO, Zhukov V, et al (2011) IPD3 Controls the Formation of Nitrogen-Fixing Symbiosomes in Pea and Medicago Spp. MPMI 24: 1333–1344

Plet J, Wasson A, Ariel F, Le Signor C, Baker D, Mathesius U, Crespi M, Frugier F (2011) MtCRE1-dependent cytokinin signaling integrates bacterial and plant cues to coordinate symbiotic nodule organogenesis in Medicago truncatula: Hormonal interactions in nodulation. The Plant Journal 65: 622–633

Reid D, Nadzieja M, Novák O, Heckmann AB, Sandal N, Stougaard J (2017) Cytokinin Biosynthesis Promotes Cortical Cell Responses during Nodule Development. Plant Physiol 175: 361–375

Roy S, Liu W, Nandety RS, Crook A, Mysore KS, Pislariu CI, Frugoli J, Dickstein R, Udvardi MK (2020) Celebrating 20 Years of Genetic Discoveries in Legume Nodulation and Symbiotic Nitrogen Fixation. Plant Cell 32: 15–41

Sakakibara H (2006) CYTOKININS: Activity, Biosynthesis, and Translocation. Annu Rev Plant Biol 57: 431–449

Sasaki T, Suzaki T, Soyano T, Kojima M, Sakakibara H, Kawaguchi M (2014) Shoot-derived cytokinins systemically regulate root nodulation. Nat Commun 5: 4983

Schiessl K, Lilley JLS, Lee T, Tamvakis I, Kohlen W, Bailey PC, Thomas A, Luptak J, Ramakrishnan K, Carpenter MD, et al (2019) NODULE INCEPTION Recruits the Lateral Root Developmental Program for Symbiotic Nodule Organogenesis in Medicago truncatula. Current Biology 29: 3657–3668.e5

Shrestha A, Zhong S, Therrien J, Huebert T, Sato S, Mun T, Andersen SU, Stougaard J, Lepage A, Niebel A, et al (2021) Lotus japonicus Nuclear Factor YA1, a nodule emergence stage-specific regulator of auxin signalling. New Phytol 229: 1535–1552

Soyano T, Kouchi H, Hirota A, Hayashi M (2013) NODULE INCEPTION Directly Targets NF-Y Subunit Genes to Regulate Essential Processes of Root Nodule Development in Lotus japonicus. PLoS Genet 9: e1003352

Tadege M, Wen J, He J, Tu H, Kwak Y, Eschstruth A, Cayrel A, Endre G, Zhao PX, Chabaud M, et al (2008) Large-scale insertional mutagenesis using the Tnt1 retrotransposon in the model legume Medicago truncatula. Plant J 54: 335–347

Tirichine L, Sandal N, Madsen LH, Radutoiu S, Albrektsen AS, Sato S, Asamizu E, Tabata S, Stougaard J (2007) A Gain-of-Function Mutation in a Cytokinin Receptor Triggers Spontaneous Root Nodule Organogenesis. Science 315: 104–107

Turner M, Nizampatnam NR, Baron M, Coppin S, Damodaran S, Adhikari S, Arunachalam SP, Yu O, Subramanian S (2013) Ectopic Expression of miR160 Results in Auxin Hypersensitivity, Cytokinin Hyposensitivity, and Inhibition of Symbiotic Nodule Development in Soybean. Plant Physiol 162: 2042–2055

van Zeijl A, Op den Camp RHM, Deinum EE, Charnikhova T, Franssen H, Op den Camp HJM, Bouwmeester H, Kohlen W, Bisseling T, Geurts R (2015) Rhizobium Lipo-chitooligosaccharide Signaling Triggers Accumulation of Cytokinins in Medicago truncatula Roots. Molecular Plant 8: 1213–1226

Vernié T, Kim J, Frances L, Ding Y, Sun J, Guan D, Niebel A, Gifford ML, de Carvalho-Niebel F, Oldroyd GED (2015) The NIN Transcription Factor Coordinates Diverse Nodulation Programs in Different Tissues of the Medicago truncatula Root. Plant Cell 27: 3410–3424

Xiao TT, Schilderink S, Moling S, Deinum EE, Kondorosi E, Franssen H, Kulikova O, Niebel A, Bisseling T (2014) Fate map of Medicago truncatula root nodules. Development 141: 3517–3528

Zürcher E, Tavor-Deslex D, Lituiev D, Enkerli K, Tarr PT, Müller B (2013) A Robust and Sensitive Synthetic Sensor to Monitor the Transcriptional Output of the Cytokinin Signaling Network in Planta. Plant Physiol 161: 1066–1075

